# What is true for plants may not be true for *Phaeodactylum tricornutum*: The case of *Vanilla planifolia* vanillin synthase (*Vp*VAN) targeted to four subcellular compartments of the diatom

**DOI:** 10.1101/2024.10.23.619700

**Authors:** Aracely Maribel Diaz-Garza, Félix Lavoie-Marchand, Natacha Merindol, Andrew Diamond, Isabel Desgagné-Penix

**Affiliations:** Department of Chemistry, Biochemistry and Physics, Université du Québec à Trois-Rivières, Trois-Rivières, Québec, Canada

**Keywords:** vanillin, diatom, organelle compartmentalization, metabolic engineering, episomal expression system

## Abstract

Vanillin is the major organoleptic compound of the vanilla flavor. Since the extraction process from its natural source (*Vanilla planifolia*) is limited by a low yield, the market demand for vanillin is met by producing it through chemical synthesis or biotechnologically using microorganisms to convert ferulic acid, lignin, and guaiacol into vanillin. Our aim was to use the diatom *Phaeodactylum tricornutum*, a photosynthetic model organism already used to produce eukaryotic proteins and high-value compounds, to characterize *V. planifolia* vanillin synthase (*Vp*VAN). This enzyme is the subject of conflicting results and was reported to be localized in the plastid of *V. planifolia* cells, which may play a role in its activity. For this purpose, we targeted *Vp*VAN linked by an *in vitro* cleavable sequence (Xa) to the enhanced green fluorescent protein (eGFP) to the cytoplasm, peroxisomes, plastid, and Golgi apparatus. We used an episomal expression system transforming *P. tricornutum* with *VpVAN:Xa:eGFP* by bacterial conjugation. Positive cell lines were selected by fluorescence and characterized by sequencing the episomes, confirming stable replication in the diatom, as well as assessing the subcellular localization in the targeted cell compartments. Strategies such as enriching GFP-positive cells by cell sorting, protein purification, and organelle isolation did not yield detectable levels of the fusion protein. A better understanding of the diatom expression system is needed to complete the characterization of this enzyme and reveal whether the subcellular localization could be a missing aspect of *Vp*VAN expression in microorganisms.

## INTRODUCTION

Natural vanilla flavor source is the orchid *Vanilla planifolia* native to Mexico and some countries from Central America [1]. The main constituent of the 200 compounds that make up vanilla flavor is vanillin (4-hydroxy-3-methoxy benzaldehyde) [2]. The low yield of extraction of vanillin from its natural source, *e. g*. one kg of vanillin requires approximately 500 kg of vanilla pods, has led to the development of alternative production using biotechnology to meet the market demand [3].

Efforts to elucidate the complete biosynthetic pathway from *V. planifolia* have led to the controversial discovery of vanillin synthase (*Vp*VAN) an enzyme with hydratase/lyase activity, which was described to catalyze the conversion of ferulic acid and its glucoside into vanillin [4]. Gallage *et al*. found *Vp*VAN to localize in the inner part of the vanilla pod with high transcripts levels in the inner epidermis [5]. Enzymatic activity was tested producing the enzyme in a coupled transcription/translation assay. Transient expression in tobacco and stable expression in barley resulted in the production of vanillyl alcohol glucoside. *Vp*VAN controversy arose from the high sequence similarity to cysteine proteases, and its identical amino acid sequence to a protein previously discovered to participate in the conversion of coumaric acid to 4-hydroxylbenzaldehyde [6]. In addition, Yang *et al*. demonstrated that the recombinant *Vp*VAN produced *in vitro*, in *Escherichia coli*, yeast or plant systems was unable to synthesize vanillin from ferulic acid. Added to the lack of tissue-specific gene expression, Yang and colleagues concluded that this enzyme cannot by itself directly convert ferulic acid to vanillin and that the pathway of vanillin biosynthesis in *V. planifolia* was yet to be determined [6].

However, heterologous production of *Vp*VAN in other plant systems, which already produce high amounts of ferulic acid and trace amounts of vanillin, such as *Capsicum frutescens* [7] and rice (*Oryza sativa* L.) calli [8], succeeded to increase the production of vanillin. This reinforced that *Vp*VAN plays a role in the production of vanillin and that there may be missing key elements in expression systems which do not produce it endogenously. Along this line, Gallage *et al*. characterized the subcellular localization of *Vp*VAN demonstrating that it localized in the chloroplast and the re-differentiated chloroplasts named phenyloplast. In addition, they isolated chloroplast and proved their biosynthetic capacity to produce vanillin from phenylalanine [5].

Even though plant systems were successful in expressing *Vp*VAN, expression of this enzyme in microorganisms has not been well established, leading to the use of alternative bacterial pathways. Ni *et al*. generated *E. coli* strains able to *de novo* synthesize vanillin from inexpensive substrates (xylose and glycerol) by introducing five genes encoding for the enzymes tyrosine ammonia-lyase, 4-coumarate 3-hydroxylase, caffeate *O*-methyltransferase, trans-feruloyl-CoA synthase (Fcs), and enoyl-CoA hydratase/aldolase (Ech) [9]. The high reactivity of the aldehyde functional group of vanillin that causes damage to the cell membrane of conventional host for metabolic engineering (i. e. *Saccharomyces cerevisiae*, and *E. coli*) limits production in microorganism cell factories [10,11]. Since accumulation of vanillin is toxic, *V. planifolia* cell strategies such as the co-expression of a UDP-glycosyltransferase has been implemented for its accumulation in yeast [12]. Also, in *S. cerevisiae* after strain optimization using a combination of engineering strategies including the blocking of the metabolic pathways responsible for the oxidation and reduction of vanillin, while the strains transformed with the *Fcs* and *Ech* from *Streptomyces*, produced up to 10.4 mg of vanillin per g of carbon source, vanillin was not detected in transgenic strains expressing *Vp*VAN [13]. A possible explanation is that compartmentalization of its production may be required to synthetize vanillin recreating the environment of phenyloplasts. Therefore, our hypothesis was that *Vp*VAN enzymatic activity depended on its subcellular localization. Thus, localizing the enzyme in the correct cellular compartment would have an impact in the production of vanillin.

*Phaeodactylum tricornutum*, a diatom which possesses a plastid, the organelle where vanillin is stored and produced in the vanilla pod [5], is already successfully used for recombinant protein expression attributed to its high growth rates and high efficiency expressing complex eukaryotic proteins [14,15]. Metabolically engineered *P. tricornutum* successfully accumulated docosahexaenoic acid (DHA) [16], produced geraniol [17], and showed an increase in biomass, lipid content, and growth [18]. In addition, the sequencing and annotation of the diatom’s genome in 2008 [19], and the development of a variety of genetic tools, have enable its use in biotechnology [20,21]. Several genetic tools have been developed, including episomal expression systems [22], TALEN technology [23], and CRISPR/Cas9 for gene knock out [24]. Promoters [25–29] and characterized subcellular localization signals were successfully identified to direct proteins to different compartments of the cell such as the plastid [30], central vacuole [31], peroxisome [32], and Golgi apparatus [33].

Therefore, the aim of this study was to identify if the subcellular localization of *Vp*VAN could influence its enzymatic activity in a microorganism cell. Our objectives were to test three subcellular compartments of the diatom cell and characterize the protein production by increasing the percentage of positive cells. We used *P. tricornutum* to express *VpVAN* linked to the enhanced green fluorescence protein (*eGFP*) with three different subcellular localization signals including bipartite signal of the chloroplast ATPase, the conserved peroxisome targeting signal 1 (PTS1), and the sequence encoding the first 23 amino acid of the xylose transferase localized in the lumen of Golgi. Subcellular localization in the targeted organelles was confirmed by confocal microscopy. Even though fluorescence was detected, *Vp*VAN recombinant protein was not. Organelle isolation for plastid constructions was used as strategy to enrich in protein fraction with *Vp*VAN as well as purification using GFP-trap without success to detect protein. This indicates that the fusion protein was produced at lower amount than the detection levels and characterization of the diatom is needed to ensure recombinant protein production.

## RESULTS

### Neither vanillin nor ferulic acid affect P. tricornutum growth at pH 8

A limitation of phenyl aldehydes heterologous production is their antimicrobial property. Therefore, prior to choosing *P. tricornutum* as a candidate organism to test *Vp*VAN’s activity, its growth was assessed under the presence of ferulic acid and vanillin. The phenolic compounds tested delayed the growth of *P. tricornutum* (Figure 1a-b). Cell morphology was assessed by light microscopy at day five (120h), revealing no observable differences between cells grown in L1 media supplemented with 1 mM of vanillin and the negative control with L1 media alone (Figure S1).

**Figure 1.**
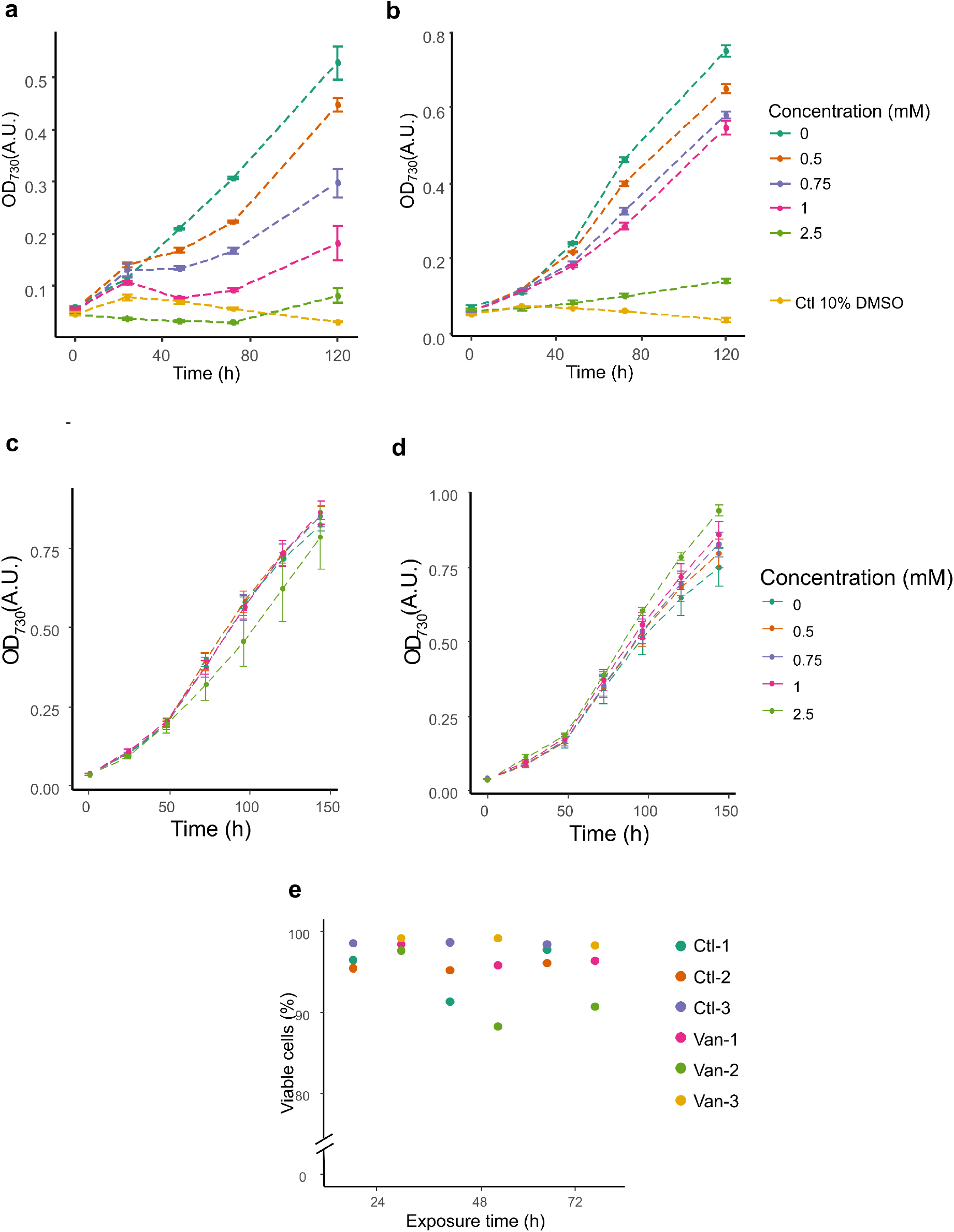
Assessing cytotoxicity of ferulic acid and vanillin to wildtype *P. tricornutum* cells. Growth of *P. tricornutum* with different concentrations of **a**) ferulic acid and **b**) vanillin, for 5 days (120h), without pH adjustment. Growth of *P. tricornutum* at pH 8 with different concentrations of **c**) ferulic acid and **d**) vanillin, for 6 days (144h). **e**. Percentage of viable cells of *P. tricornutum* incubated with 2.5 mM of vanillin for 24, 48 and 72h. n= 3; Ctl: L1 media alone; Van: 2.5 mM of vanillin; numbers indicate biological replicates.

After adjusting pH of the media to 8, neither vanillin nor ferulic acid had a negative effect in *P. tricornutum* growth (Figure 1c-d). In fact, growth was slightly higher at 144h of cells cultured with 2.5 mM of vanillin (Figure 1d). Cell viability was also assessed in the presence of 2.5 mM of vanillin at pH 8 using propidium iodide (PI), showing no cytotoxic effect at 24, 48, or 72h (Figure 1e).

### Selection and characterization of P. tricornutum exconjugants

To target and assess *Vp*VAN localization in specific compartments, several constructions were assembled, *VpVAN* was linked to *eGFP* through the cleavable peptide by factor Xa. To target the chloroplast, *VpVAN* was preceded by a signal specific to the plastid stroma in pCS*Vp*VAN:eGFP (Figure 2). To avoid hampering the normal function of *P. tricornutum* plastid, we assembled alternative constructions p*GoVpVAN:eGFP* and p*PeVpVAN:eGFP* to direct the enzyme to Golgi and peroxisome, respectively, as well as one construction without any signal to target the cytosol (p*CyVpVAN:eGFP*). The signal peptide to target the Golgi was placed before *VpVAN*, while it was placed after e*GFP* for localization in peroxisomes.

**Figure 2.**
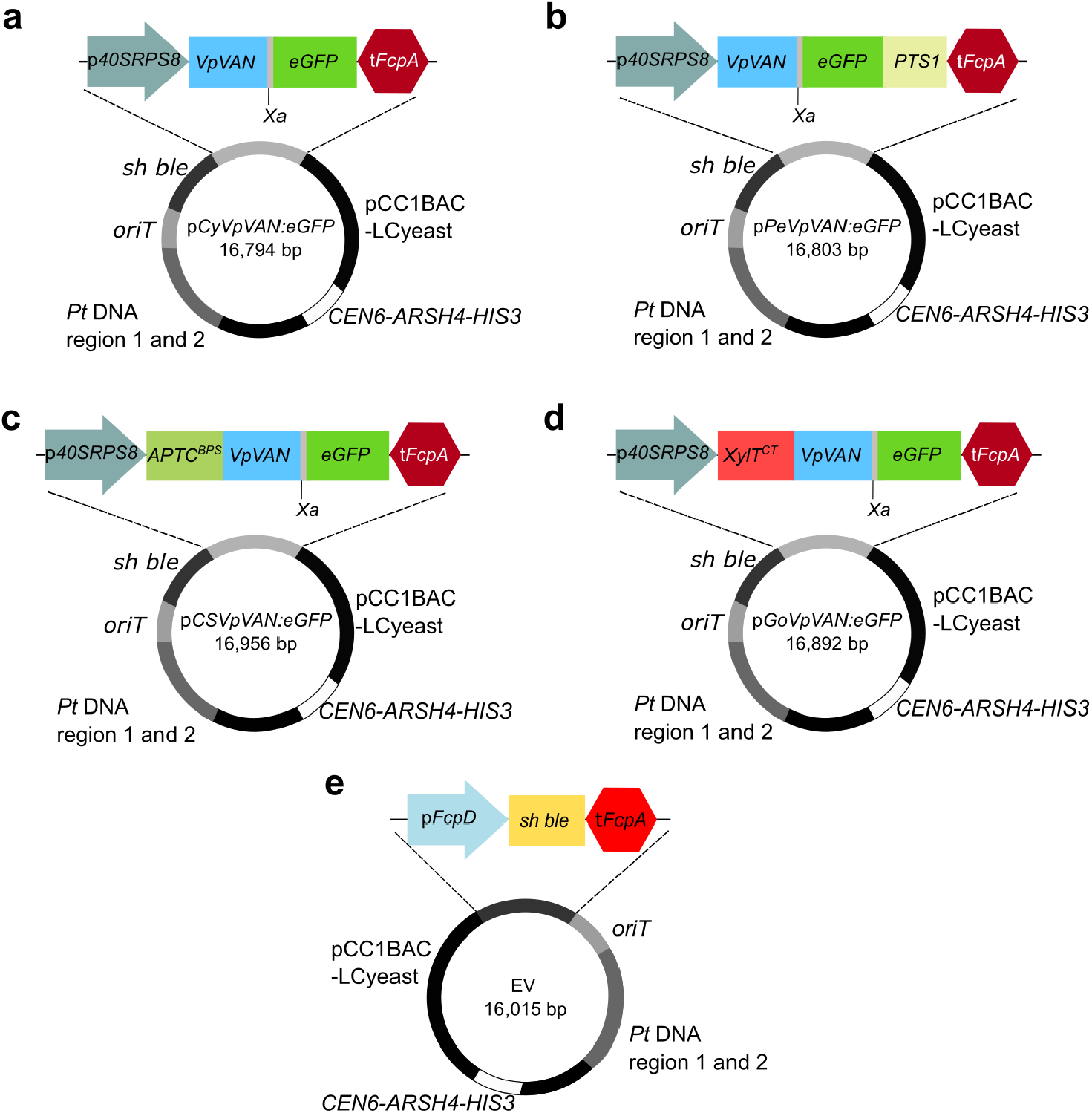
Stable episome propagation without mutations in the expression cassettes with localization signals for three different organelles and the cytosol. **a**. Cytosolic construction of *VpVAN:Xa:eGFP* without any localization signal; **b**) peroxisome construction harboring the peroxisomal targeting signal 1 (*PST1*); **c**) plastid construction with the bipartite signal of the chloroplast ATPase gamma subunit (*ATPC*^BPS^); **d**) β1,2-xylosyltransferase cytosolic tail and transmembrane domain (CT) for medial Golgi (*XylT*^CT^). *Vp*VAN expression cassette is antisense from *sh ble*. **e**. Empty vector strain harboring the pPtGE30 plasmid. Distances are not at scale.

Following transformation and a two-week selection in *P. tricornutum in* ½L1 agar plates with zeocin, at least 36 colonies of each construction were screened for fluorescence using an epifluorescence microscope (Table 1). The percentage of fluorescent colonies for each construction is presented in Table 1. p*CSVpVAN:eGFP* transformants exhibited the lowest percentage of positive colonies with 53.7%. The peroxisome construction had the highest percentage with 80.5% positive colonies. Three different positive colonies per construction were picked for further analyses. To confirm the integrity of the gene of interest, whole plasmid sequencing was performed, revealing no mutations in the expression cassettes in any of the clones (Figure 2).

**Table 1.** Fluorescent colonies per construction.

| Construction | Percentage of fluorescent colonies |
| --- | --- |
| pCyVpVAN:eGFP | 24/36 (66.7%) |
| pPeVpVAN:eGFP | 29/36 (80.5%) |
| pCSVpVAN:eGFP | 29/54 (53.7%) |
| pGoVpVAN:eGFP | 25/36 (69.4%) |

### Successful localization of the recombinant proteins in four different cell compartments

The assessment of the subcellular localization was done for three different transformants for each construction by confocal microscopy. Fluorescent signal from the clones carrying *VpVAN:Xa:eGFP* was compared to the empty vector (EV) strain as negative control (Figure 3a). Clones were only considered positive for protein production if the fluorescent pattern was different from the EV strain, including higher signal intensity. Localization in the cytosol, visible as a uniform signal throughout the cell, was confirmed in Cy*Vp*VAN:eGFP clones (Figure 3b). In the case of Pe*Vp*VAN:eGFP and Pe:eGFP clones, a punctuated pattern of three to five dots surrounding the plastid was observed (Figure 3c). The signal for these constructs was also detected in the cytosol.

**Figure 3.**
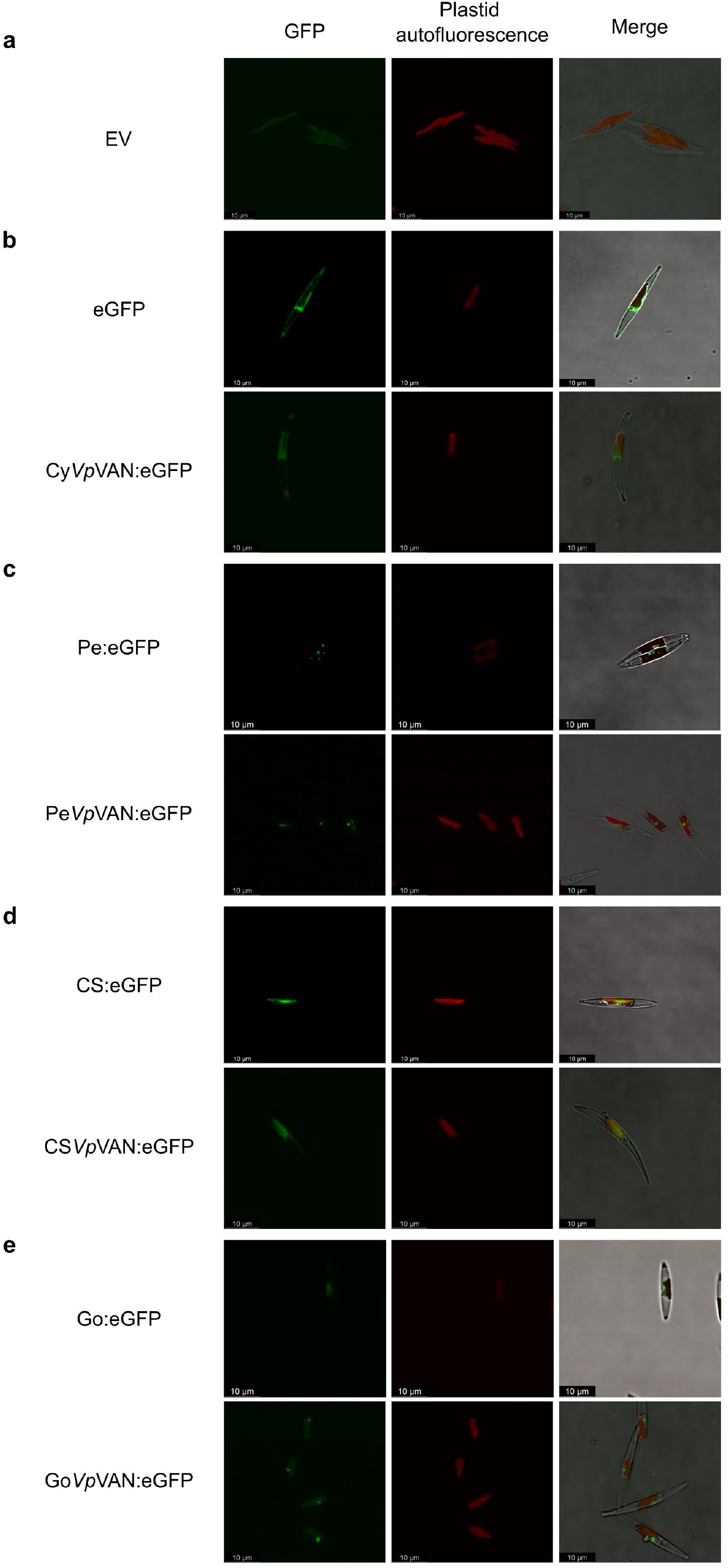
The fusion protein VpVAN:Xa:eGFP localizes in the targeted cell compartment of *P. tricornutum*. **a**. EV: empty vector strain. **b**. cytosolic constructions: eGFP and Cy*Vp*VAN:eGFP. **c**. Pe:eGFP and Pe:*Vp*VAN:eGFP clones. **d**. plastid stroma clones with the bipartite signal of ATPC linked to eGFP alone (CS:eGFP) and the fusion protein (CS*Vp*VAN:eGFP). **e**. Clones with Golgi retention signal: Go:eGFP and Go*Vp*VAN:eGFP. Merge images represent the combination with plastid autofluorescence, GFP channel and bright field. Scale bars 10 μm.

Interestingly, differences between controls and clones with *Vp*VAN constructions were observed for plastid and Golgi subcellular localization. Co-localization of the recombinant protein and plastid autofluorescence was observed in CS*Vp*VAN clones (Figure 3d), as well as aggregation of proteins seen as high intensity dots together with cytosolic fluorescence. Clones harboring the Golgi retention signal, with and without *Vp*VAN, presented a characteristic “C”-shape in the middle of the plastid (Figure 3e). In addition, Go:eGFP clones also presented fluorescence in the endoplasmic reticulum (ER). This pattern was not observed in Go*Vp*VAN:eGFP clones, which instead had overlapping fluorescence with the plastid. The discrepancy between control and the localization of *Vp*VAN clones could be attributed to the amino acids located after the predicted transmembrane domain (TMD) from XylT. *In silico* analysis predicted the three first amino acids of eGFP as part of the TMD of Go:eGFP, not present in either the full length XylT or Go*Vp*VAN:eGFP.

Since most of the localization signals are in the N-terminal of the fusion protein and sequencing results confirmed intact cassettes, we could deduce that the fusion protein is present in the positive clones. However, the main cause of *Vp*VAN controversy is the inability to reproduce vanillin synthesis by itself. Therefore, it is vital that we confirm the production of the full fusion protein in our desired cellular compartments.

### Enrichment in fluorescent cells, protein purification, and plastid isolation failed as strategies to detect heterologous protein by western blot in positive clones

We aimed to enrich in fluorescent cells by FACS with the purpose of increasing the percentage of cells in one clone that produce the fusion protein. We used an empty vector strain to gate the GFP+ cells (Figure 4a). However, after sorting GFP+ cells and growing them for four weeks we were not able to obtain high percentage of fluorescent cells (Figure 4b-e).

**Figure 4.**
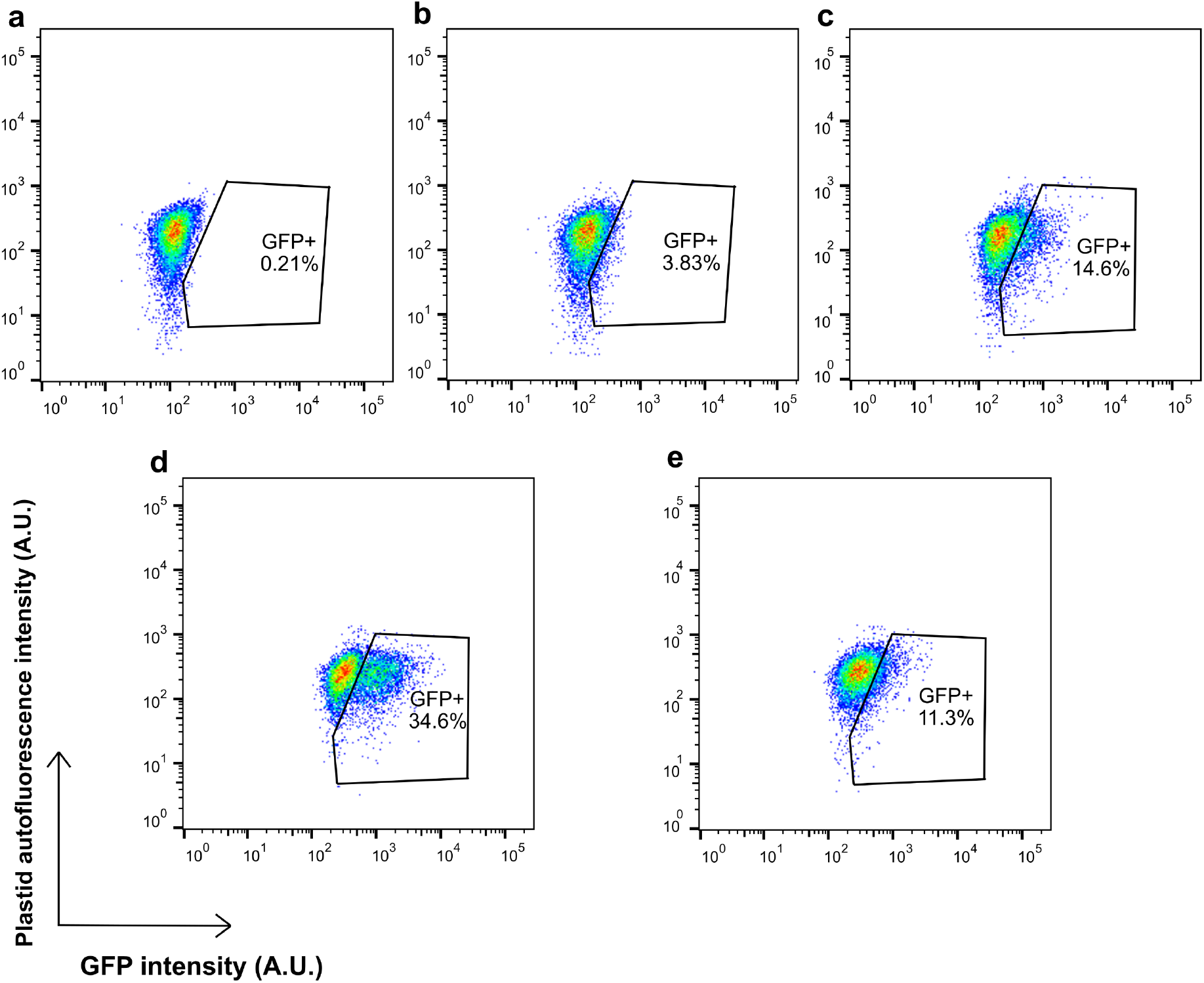
Enrichment for fluorescent cells by FACS fails to yield a high percentage of fluorescent cells. A single cell line is presented per targeted cellular compartment, four weeks after sorting. **a**. empty vector (EV); **b**. Cy*Vp*VAN:eGFP; **c**. Pe*Vp*VAN:eGFP; **d**. CS*Vp*VAN:eGFP; **e**. Go*Vp*VAN:eGFP.

*Vp*VAN production was not detected in any of the transformed cell lines by western blot, mainly attributed to the low percentage of fluorescent cells. Since we were not able to increase the number of fluorescent cells in the population by cell sorting, we decided to enrich the protein fraction to detect the protein by western blot. We used two different strategies: plastid isolation for the CS*Vp*VAN clones and protein purification by GFP-trap for Cy*Vp*VAN clones. Unlike for peroxisomes and Golgi, plastid isolation protocol for *P. tricornutum* is available in the literature. Besides, the CS*Vp*VAN:eGFP had the highest percentage of fluorescent cells. Purification using GFP-trap was done only for the cytosolic clones since it is the only one where the protein is not compartmentalized in another membranous structure. However, western blots failed to reveal bands specific to *Vp*VAN, thus protein production of the fusion protein was not detected in any of the tested clones (Figure 5). The controls of protein transfer to the membrane are present in Figure S2. We succeeded to purify eGFP:T2A and eGFP:T2A:mCherry from a pDMi8 cell line as a positive control (Figure 5a), the detected bands at 36kDa and 58 kDa correspond to the cleaved (eGFP:T2A) and uncleaved (eGFP:T2A:mCherry). The percentage of GFP+ cells for the control was 35.2 %.

**Figure 5.**
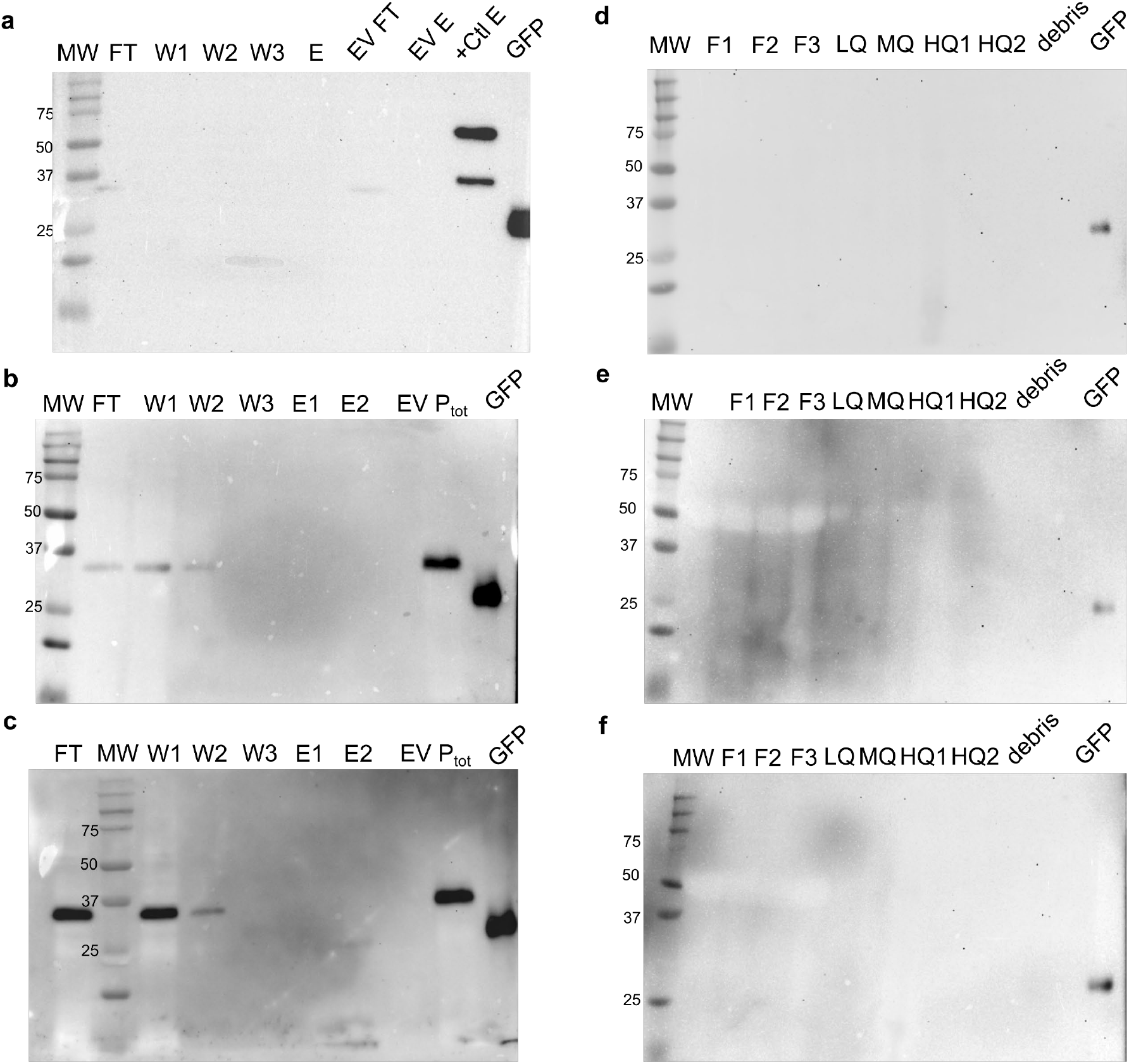
Enrichment of protein fraction and purification failed to increase fusion protein to detectable levels. **a**-**c**. Western blot of protein extracts from three different clones with cytosol constructions after purification using GFP-trap. **d**-**f**. Western blot using protein extract of plastid purification of three different clones harboring the plastid construction. MW: molecular weight marker; FT: flow through; W1-3: washes; E1-2: elution fraction; P_tot_: total protein in the soluble fraction; +Ctl: eGFP:T2A:mCherry clone; F1: total protein extract; F2: thylakoids fragments and mitochondria; F3: plastids; LQ: low quality plastids; MQ: medium quality plastids.

## DISCUSSION

To first assess if *P. tricornutum* was a suitable platform to produce vanillin we observed the diatom growth in 2.5 mM of vanillin. Vanillin has been reported to be toxic for *E. coli, Lactobacillus plantarum* and *Listeria innocua*, since it is a membrane-active compound able to dissipate ion gradients and inhibits respiration [34]. *P. tricorntum* growth was delayed when cultured with ferulic acid and vanillin without adjusting the pH of the media. However, after we adjusted the pH to 8, we did not observe any delay in growth, we even observed a higher growth when culturing *P. tricornutum* at 2.5 mM of vanillin. Studies show that vanillin is a transient intermediate of ferulic acid degradation and can be used as carbon source by 42 strains of Actinobacteria [35]. It has been proven that by increasing the pH of the media, ferulic acid occurs in its anionic form and can be transported better into the cell with less membrane damage increasing both, ferulic acid consumption rate and vanillin yield in an engineered *E. coli* strain harboring *Ech* and *Fcs* [36]. We validated the fitness of the cells at 2.5mM of vanillin pH 8 by assessing membrane integrity. We did not observe significant cytotoxic effect after 24, 48, or 72h adding vanillin to the media. While our results support the use of *P. tricornutum* as a platform to produce phenylpropanoids, they also indicate that metabolic optimization is needed since phenyl compounds could be metabolized by the cell.

We constructed four different expression cassettes to target *Vp*VAN to cytosol, peroxisomes, plastid stroma, and Golgi apparatus. While in *V. planifolia* this enzyme is localized in the plastid, we decided to target the production of vanillin to other cellular compartments in *P. tricornutum* since previous studies have shown that the accumulation of phenylpropanoids in a plastid can cause a re-differentiation, stopping the production of chlorophyll and thus the photosynthesis [37]. The Golgi apparatus is an organelle of interest since glycosyltransferases [38] reside in this compartment. Therefore, targeting the production of vanillin to Golgi could yield production of vanillin glycoside, which can be accumulated in the cell without presenting toxicity. Peroxisomes have a single-layer membrane where low molecular weight compounds can cross passively or by protein channels [39]. These organelles are in charge of detoxification which means they can sequester and handle molecules that are toxic for the rest of the cell, such as vanillin [39]. In addition, a re-evaluation of vanillin biosynthetic machinery has been proposed since Gallage *et al*. only confirmed the subcellular localization of *Vp*VAN [5]. It could be possible to have cytosolic and ER-anchored enzymes that were also isolated with the chloroplast tested as co-isolation of the fragments of other organelles with chloroplasts is common to happen [40].

After selecting three positive colonies of each construction, we verified the integrity of the episomes by sequencing. While plasmid re-arrangements have been described to happen in *P. tricornutum* [41], our results are consistent with previous reports showing stable plasmid propagation in the diatom [21,42].

The subcellular localization patterns matched what has been previously described for the localization signals used in this study [30,32,33]. However, we observed differences in eGFP signal comparing clones carrying *VpVAN* and the control clones with eGFP linked to the localization signals. The CS*Vp*VAN:eGFP clones presented besides the chloroplast localized fluorescent signal, aggregation of proteins observed as high intensity dots and cytosolic fluorescence not seen in the control without *Vp*VAN (p*CSeGFP*). This distinct fluorescent pattern could be caused by a cleavage of the Xa peptide by native *P. tricornutum* peptidases, as it has been hypothesized by Messaabi and colleagues producing free eGFP, localizing in the cytosol [43].

Optimization of protein compartmentalization may be necessary for the peroxisome and Golgi localizations. Clones harboring PTS1 presented cytosolic fluorescence most likely due to the presence of the recombinant protein prior to its import. An enhanced PTS1 sequence in *S. cerevisiae* has been reported to increase the rate of cytosolic protein import by the addition of the linker LGRGRR, which has a higher affinity to the import receptor [44]. A similar strategy could be implemented for the diatom to effectively compartmentalize the enzymatic activity into the peroxisome. There is evidence in other organisms that a single TMD, arginine-based motifs residues used as sorting sequences, and protein conformation are responsible in the localization of glycosyltransferases within Golgi [45]. Moreover, it has been shown that Golgi-resident enzymes possess multiple mechanisms to effectively be retained in this compartment, which can be complementary or additive [46]. Therefore, a characterization is needed to identify the elements in XylT sequence responsible of targeting this enzyme to Golgi and obtain reproducible results using different proteins. CTS regions of glycosyltransferases from other organisms contain a luminal stem region separating the catalytic domain from the TMD that plays a role in sub-localization and oligomerization of the protein, in addition to the cytosolic tail and a single short TMD [47,48]. Identification of the minimal stem region necessary to retain enzymes in Golgi could replace the use of the full amino acid sequence to target proteins to this compartment. Alternatively, the fragment tested in this study could provide a shared localization in Golgi and ER, which may be beneficial for accumulation of vanillin glycoside since both compartments harbor glycosyltransferases [49,50].

Detection of the *Vp*VAN:Xa:eGFP protein by western blot was not possible in this study. A plausible cause of the lack of signal could be very low amounts of produced protein that despite purification was below the detection limits of our assay. This led us to use different strategies to increase the protein fraction of our protein of interest including, purification, organelle enrichment and increasing the fraction of GFP+ cells by FACS. However, these three strategies failed to yield detectable levels of our protein of interest. In addition, we obtained low percentages of GFP+ cells after growing the sorted cells from our positive clones for four weeks. These results indicate that we selected more than one phenotype when sorting the cells and/or *P. tricornutum* regulated the production of the fusion protein in days post-sorting. Our results are consistent with George and colleagues who were not able to enrich in phenotypic populations by cell sorting of *P. tricornutum* cells episomally expressing *mVenus* [17]. Studies done in heterologous subpopulations in cell lines expressing *mVenus* integrated into the genome of *P. tricornutum* suggest that differential DNA methylation patterns and histone modification (H3K4me2) are not responsible for transgene silencing [51]. In addition, in another study, we reported differences in plasmid copy number, a large deletion, and co-expression of a putative zeocin resistant gene with the transgene, as factors contributing to the formation of subpopulations in a single clonally propagated episomal cells line of *P. tricornutum* [52].

The decrease in heterologous protein production may be specific to cell lines harboring *Vp*VAN, as a response for the high production of an enzyme with high amino acid sequence homology to a cysteine protease. Identification of protein degradation mechanisms in *P. tricornutum*, by using protease or proteasome inhibitors, could provide insight into the accumulation of heterologous proteins, although this could affect *Vp*VAN cysteine protease function. Additionally, this information could lead to the engineering of strains with lower protein degradation mechanisms. Similar work has been done in yeast strains, generating mutant variants with disrupted protease genes, efficiently increasing heterologous protein accumulation [53,54].

An alternative hypothesis regarding the low accumulation of *Vp*VAN is that the trace amount of ferulate in *P. tricornutum* [55] could be used by the enzyme to produce vanillin, and the cell would in turn downregulate its production. The diatom cells could be adapting to reduce the production of a toxic enzyme or product, as it has been suggested in our previous study, where re-arrangements occur in high frequency in an episome harboring *Ech* and *Fcs* genes, able to convert ferulic acid into vanillin [41]. Diatoms have evolved to rapidly adjust to a changing environment, therefore mechanisms lighten the metabolic burden of producing a toxic enzyme may be in place to decrease the synthesis of *Vp*VAN. Better understanding of dynamics of *P. tricornutum* expression system is needed to successfully characterize possibly toxic proteins such as *Vp*VAN.

## CONCLUSIONS

Synthesis of vanillin from ferulic acid in one-step using *Vp*VAN remains a challenge, since many characteristics of this enzyme are still unknown. However, by establishing requirements for the expression of *Vp*VAN in microorganisms, such as targeting a specific compartment of the cell, we could contribute to answer questions about its activity. In this study we were able to introduce *VpVAN*:*Xa:eGFP* in an episome by bacterial conjugation into *P. tricornutum*. Plasmid rescue and sequencing showed that episomes remained extrachromosomal and were stably propagated. Besides, we confirmed that previously characterized localization signals of the diatom effectively directed our fusion protein into the plastid, peroxisome, and Golgi. However, we also observed that optimization of this sequences to increase the rate of transport of the desired protein may be needed to avoid pre-import enzymatic activity. Finally, we were not able to detect either truncated or full protein by western blot. Enrichment strategies used by previous studies failed to yield detectable levels of the fusion protein, which questions if this effect is due to the alternative enzymatic activity proposed of *Vp*VAN as cysteine protease exhibiting toxicity for the diatom cell. These results strongly suggest that *P. tricornutum* expression system needs to be studied in more details to be able to characterize “toxic” enzymes.

## MATERIALS AND METHODS

### Microbial Strains and Growth Conditions

*Escherichia coli* (Epi300, Epicenter) was grown in Luria Broth (LB) supplemented with appropriate antibiotics (gentamicin (20 mg L^-1^) or chloramphenicol (15 mg L^-1^) and gentamicin (20 mg L^-1^)). *Phaeodactylum tricornutum* (CCAP 1055/1, Culture Collection of Algae and Protozoa) was grown in modified L1 medium without silica at 18 °C under cool white fluorescent lights (75 μE m^−2^ s^−1^) and a photoperiod of 16 h light:8 h dark with an agitation of 130 rpm for liquid cultures [41].

### Cytotoxicity assays of P. tricornutum with ferulic acid and vanillin

To observe the effect of phenylpropanoids in the diatom’s growth, 2.0 × 10^6^ *P. tricornutum* cells were cultured in 200 μL in a 96-well plate with L1 media for 5 days with 0, 0.5, 0.75, 1, 2 mM of either ferulic acid or vanillin, and its growth was assessed by measuring each 24h the optical density at 730 nm. The experiment was repeated adjusting the pH of the media to 8 and assessing growth for 6 days. Cells grown in 1mM of vanillin were assessed for changes in their morphology at day five by light microscopy (OMAX, USA) using the 40X objective. Finally, the percentage of viable cells of cultures with 2.5 mM of vanillin after 24, 48, and 72 hours of treatment was measured by incubating for 10 min with propidium iodide (PI) final concentration of 3 μg/mL. PI emission was measured by BD FACS Melody (BD Biosciences, La Jolla, CA, USA) in the ECD channel (610/20 nm). The percentages of live cells were calculated relative to the untreated control wells.

### Plasmid construction and transformation of P. tricornutum by bacterial conjugation from E. coli cells

Plasmids were constructed by Gibson Assembly using the NEBuilder® HiFi DNA Assembly Bundle for Large Fragments (New England Biolabs, Canada). All fragments used in the assemblies were amplified by PCR using PrimeSTAR GXL DNA Polymerase (Takara Bio, Japan) following the manufacturer’s protocol. Episomes were completed by replacing the coding sequence in the expression cassette of pDMi7 [41] containing the *40SRPS8* (40S ribosomal protein S8) promoter and the *FcpA* terminator, with the backbone of pPtGE30 harboring the *CAH* region (*CEN6-ARSH4-HIS3*) and the *sh ble* gene that confers resistance to zeocin. All expression cassettes consisted in *VpVAN* without the predicted signal peptide linked to the enhanced green fluorescent protein gene (*eGFP*) by the factor Xa cleavable peptide (*Xa*). Depending on the targeted cellular compartment *P. tricornutum* localization signals were added to each construction: cytosol without any signal, peroxisome construction harboring the peroxisomal targeting signal 1 (PST1) tripeptide SKL, plastid construction with the bipartite signal of the plastid ATP synthase gamma chain (Phatr3_J20657) to direct the recombinant protein to the stroma, and Golgi apparatus with the N-terminal (31 amino acids) of the β1,2-xylosyltransferase (Phatr3_J45496). The latest contains a predicted transmembrane domain as well as and a triple arginine motif (RRR) in the cytosol tail. In addition, four constructs harboring the localization signals and *eGFP* were assembled as controls for subcellular localization (Figure S3). The empty vector used in this study was the plasmid pPtGE30 [21], harboring no expression cassette besides *sh ble*.

Sequences were synthesized and optimized for *P. tricornutum* codon usage by Bio Basic (Markham, Ontario, Canada). All primers used in the assembly are listed in Table S1. DNA sequences from this study are available in Table S2.

Successful cloning and transformation of all four constructions was confirmed by screening 10 chloramphenicol resistant *E. coli* NEB 10-beta colonies by colony PCR. A single colony for each construction was chosen for plasmid extraction and the integrity of the episomes was verified by Sanger sequencing (Génome Québec). Plasmids with intact expression cassettes were transformed into *E. coli* Epi300 pTA-MOB, since this strain harbors the plasmid that allows to do conjugation with *P. tricornutum*.

Episomes were transformed into wild type *P. tricornutum* by bacterial conjugation as previously described [41]. Briefly, 250 μL of *P. tricornutum* culture, concentration of 1.0 × 10^8^ cells mL^-1^, were plated in ½L1 agar plates and grown under the conditions mentioned above for 4 days. Prior to transformation, cells were harvested by scraping the plates and adding 1 mL of L1 media, cell concentration was then adjusted to 5.0 × 10^8^ cells mL^-1^. A culture of 25 mL of *E. coli* EPI300 pTA-MOB containing either the empty vector or pDMi8 plasmids was grown at 37°C and 220 rpm until reaching an OD^600^ of 0.7, then centrifuged at 3000 g for 10 min and resuspended in 250 μL of SOC media. To initiate the conjugation, 200 μL of *P. tricornutum* cells were mixed with 200 μL *E. coli* cells, plated in ½L1 5% LB agar plates, and incubated at 30°C for two hours in the dark. After conjugation, plates were kept at standard growth conditions for *P. tricornutum* for the recovery period of 2 days. Cells were collected by scraping with 1 mL of L1 media and plated in selective ½L1 agar plates with zeocin 50 μg/mL. Colonies appeared after 10 to 14 days of growth at 18°C and photoperiod of 16 h light:8 h dark.

### Screening positive exconjugants by fluorescence microscopy

Two weeks following the conjugation colonies were screened by eGFP fluorescence using a Fluorescent Stereo Microscope Leica M165 FC with GFP filter for eGFP fluorescence detection. Colonies were observed with a magnification of 80 to 120x. Three positive colonies for each construction were selected to start 25 mL cultures.

### Episome rescue, whole plasmid sequencing and in silico analysis

Episome isolation from P. tricornutum was conducted as described in Diamond et al. [41], using Large Plasmid Mini Kit (Geneaid Biotech Ltd., Taiwan), with approximately 1 × 10^8^ cells of P. tricornutum. Plasmid extracts were used to transformed NEB 10-beta chemically competent E. coli cells following the manufacturer’s protocol. After overnight incubation of the plates a single colony was used to inoculate 10 mL LB cultures to proceed for miniprep using Large Plasmid Mini Kit (Geneaid Biotech Ltd., Taiwan). Plasmids were sequenced by CCIB DNA Core (Massachusetts General Hospital, United States of America) through their Next-Generation sequencing Illumina MiSeq platform.

Amino acid sequences obtained from in silico translation of the sequenced episomes targeting Golgi were analyzed using TMHMM 2.0 as well as the full XylT sequence to predict the transmembrane domains [56].

### Assessing subcellular localization by confocal microscopy

Live cell images of the five-day-old culture were captured using a Leica SP8 confocal laser microscope (Leica, Wetzlar, Germany) with an HCX PL APO 60×/1.25–0.75 Oil CS objective. The excitation of eGFP and chlorophyll fluorescence occurred at 488 nm with a 65 mW argon laser. eGFP fluorescence emission was detected between 499nm and 531nm, whereas chlorophyll fluorescence was detected at a bandwidth of 640nm - 710nm. Bright-field light microscopy images were also taken. The images were analyzed using LAS X software (version) (Leica).

### Fluorescence-activated cell sorting (FACS)

The BD FACS Melody (BD Biosciences, La Jolla, CA, USA) equipped with blue (488 nm), red (640 nm) and violet (405 nm) lasers was used to analyze and sort *P. tricornutum* transconjugants according to eGFP production as described by Diamond *et al*.[41]. Five-day-old *P. tricornutum* cultures were filtered with Falcon™ Cell Strainers (Fisher Scientific, USA) prior to sorting. Events were acquired at a fixed flow rate and at least 10,000 events were analyzed. Cells were gated according to FSC-A (forward scatter area) and SSC-A (side scatter area) parameters and doublets were excluded according to further gating on homogeneous FSC-H (height) *vs*. FSC-W (width) and SSC-H *vs*. SSC-W populations. Chloroplast autofluorescence was gated in the PerCP channel (700/54 nm, 665 LP). Cells with non-specific autofluorescence detected in the PE channel (582/15 filter, 582 long pass filter mirror – LP) were excluded from sorting. eGFP was further analyzed on the 527/32 nm band-pass filter channel. Sorting was set on purity parameter. Sorted cells were collected in 1.5 mL microcentrifuge tubes containing 250 μL of L1 media with ampicillin (100 μg/mL) and zeocin (50 μg/mL). Cells were centrifuged for 10 min at 3,500 g, supernatant was removed to avoid toxicity from the FACS sheath fluid, and fresh L1 media with antibiotics was added. Cultures were grown for two weeks and used as inoculum for 50 mL cultures. Data analysis was conducted with BD FlowJo version 10 software (BD Biosciences, La Jolla, CA, USA, 2020).

### Protein extraction and purification

Protein was extracted using 20 mL seven-days-old cultures as described by Fantino *et al*. (2024). Cells were harvest by centrifugation at 3,500 x*g* for 10 min at 4°C. Pellets were weighted and 500μL of lysis buffer (51.4 mM Tris pH 8, 0.75 mM SDS, 10 % Glycerol, 0.02 mM EDTA, 10 mM PMFS and 2 μL of protease inhibitor cocktail) per 200 mg of wet biomass was added. Cells were lysed by sonication with 35% of amplitude and 6 pulses of 30 s with a rest period of 30 s between pulses using Fisherbrand Model 505 Sonic Dismembrator (Thermo Fisher Scientific). The soluble fraction was separated by 30 min of centrifugation at 20,000 x*g*. The supernatant of clones harboring the cytosol construction was used for purification with GFP-trap beads following the manufacturer protocol, keeping aliquots of each fraction for posterior analyses. Protein quantification was performed using the RC DC Protein Assay Kit I (Bio-Rad).

### Plastid isolation

Plastid isolation from clones carrying the construction targeting this organelle was performed according to the protocol of Schober *et al*. [58] with the modification of using sonication as mentioned above to lyse the cells. After cell disruption, intact cells and debris were removed by centrifugation at 500 x*g*. The supernatant containing mainly plastids and mitochondria was centrifuged at 2,000 *xg* to concentrate the plastids. The supernatant was discarded, and the pellet was resuspended in isolation buffer (0.5 M Sorbitol, 50 mM HEPES-KOH pH 8, 6 Mm EDTA, 5 mM MgCl_2_, 10 mM KCl, 1mM MnCl_2_, 1% polyvinylpyrrolidone, 0.5% BSA, and 0.1% L-cysteine). High quality plastids were isolated by applying the plastid-containing solution into a layered Percoll density gradient and centrifuged at 10,000 *x g*. Plastid were washed by resuspending them in washing buffer (0.5 M Sorbitol, 30 mM HEPES-KOH pH 8, 6 mM EDTA, 5 mM MgCl_2_, 10 mM KCl, 1mM MnCl_2_, and 1% polyvinylpyrrolidone) and centrifugated at 4,000 x*g*. Proteins were extracted from the final plastid fraction by freeze and thaw (using liquid nitrogen), and resuspending on protein extraction buffer containing 10% glycerol, 5mM EDTA, 10 mM DTT, and 100 mM HEPES (pH 7.2).

### Western blot

Proteins were detected by western blot, loading 50 μg for soluble fraction. SDS-PAGE of 12% polyacrylamide. Regarding the washing and elution fractions, 25 μL of sample were loaded since the amount protein was either too low or protein quantification was not possible due to interaction with the reagents of the elution buffer. Also, a 3 μL of positive control for western blot (purified GFP) were loaded in the gel. Protein transfer to a 0.2 μm PVDF membrane was done by semi-wet transfer at constant amperage of 1 A for 30 min and maximum voltage of 25 V. The membrane was blocked using 3% BSA in Tris-buffered saline, 0.1 % Tween 20 (TBST) solution for one hour at room temperature. Afterwards it was incubated overnight with anti-GFP antibody (CLH106AP) purchased from Cedarlane (Ontario, Canada). Then, the membrane was washed using TBST and incubated for 1 h in a 1:20,000 dilution in 5 % milk of Immun-Star Goat Anti-Mouse (GAM)-HRP conjugate from Bio-Rad (Ontario Canada). Three washes were done, and protein detection was performed using Clarity Max Western ECL Substrate-Luminol solution from Bio-Rad. Chemiluminescence detection and Ponceau S stained (Glacial Acetic Acid 5 % *v*/v, Ponceau Red dye 0.1 % m/v) of the blots were visualized using ChemiDoc Imaging System with Image Lab™ Software (Bio-Rad). The molecular weights of the proteins corresponding to the detected bands were confirmed with protein markers (Precision Plus Protein Dual Color Standards).

## DATA AVAILABILITY STATEMENT

The original contributions presented in the study are included in the article and supplementary information, further inquiries can be directed to the corresponding author.

## ACKNOWLEDGMENTS

Warm thanks to Elisa Fantino for the lab training in molecular biology techniques. This research was funded by Canada Research Chair on plant specialized metabolism Award No CRC-2018-00137 to I.D-P. Thanks are extended to the Canadian taxpayers and to the Canadian government for supporting the Canada Research Chairs Program. Additional support in the form of scholarship to A.M.D-G. was provided by Mitacs-Acceleration program grant #IT12310.

## AUTHOR CONTRIBUTIONS

ID-P, AD, and AMD-G conceived the idea of the project; FL-M, NM, and AMD-G. designed methodology and performed experiments; AMD-G wrote the first draft and created the figures using Inkscape; All authors have read and corrected the manuscript.

