## Supplementary information for "What is true for plants may not be true for *Phaeodactylum tricornutum*: The case of *Vanilla planifolia* vanillin synthase (*Vp*VAN) targeted to four subcellular compartments of the diatom"

**SUPPLEMENTARY DATA**

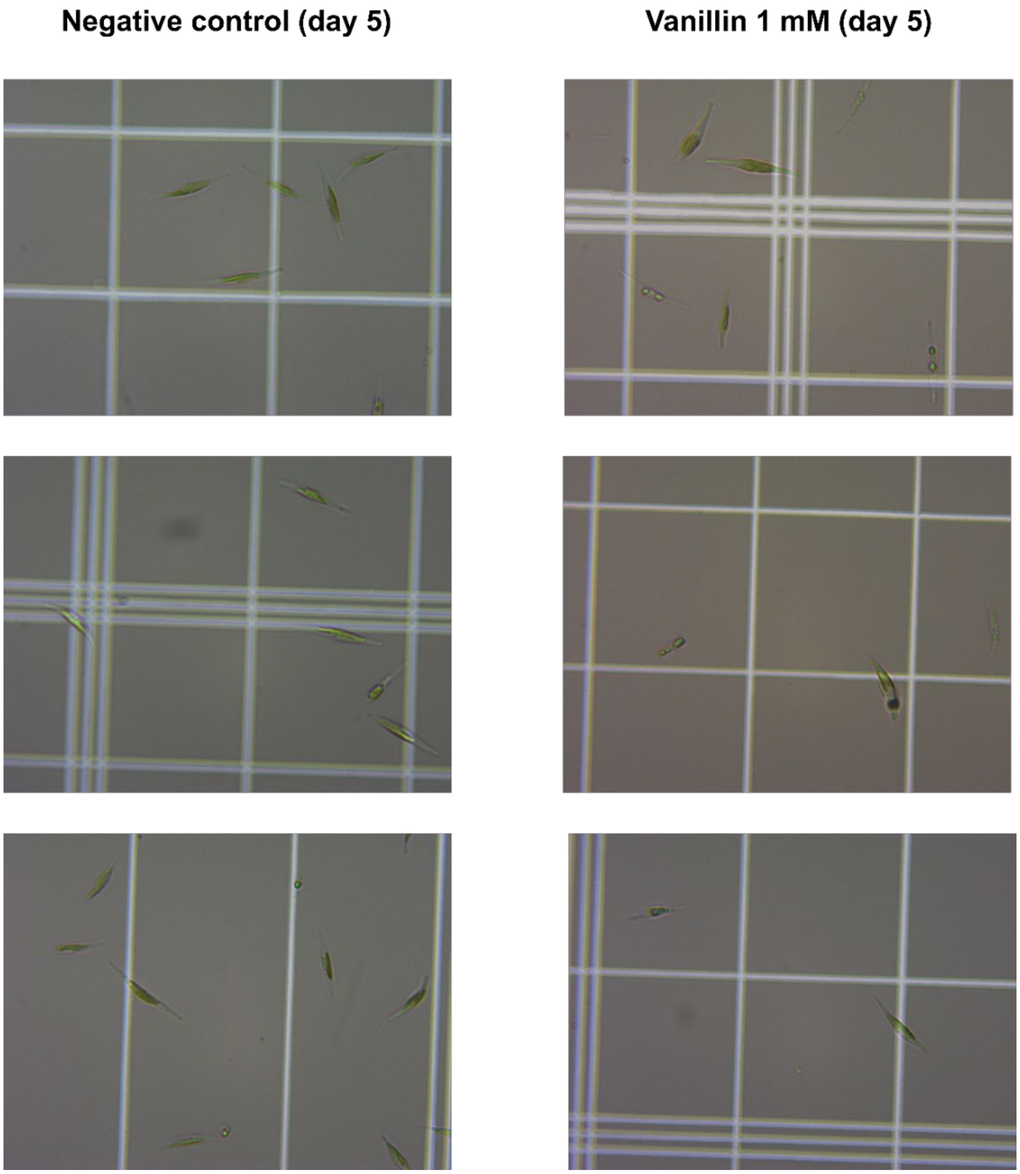


**Figure S1. The morphology of *P. tricornutum* cells treated with 1 mM of vanillin is similar to untreated cells at day five.** Three biological replicates of the wild-type strain are shown for the non-treated (left panels) and treated cells with vanillin (right panels). Microscopy images were obtained using the 40x objective.

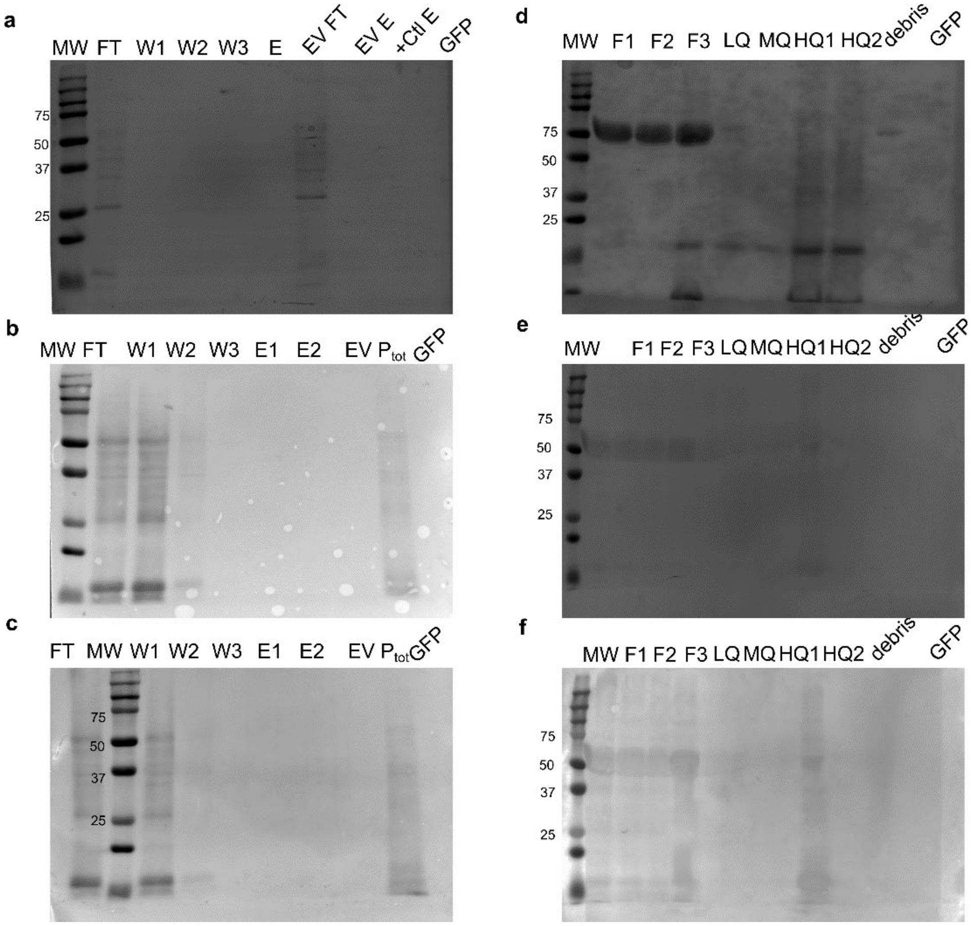


**Figure S2. Proteins transferred to the blotting membrane.** Red ponceau staining of membranes used for western blot: (**a-c**) protein extracts from three different clones with cytosol constructions after purification using GFP-trap; (**d-f**) protein extract of plastid purification of three different clones harboring the plastid construction. MW: molecular weight marker; FT: flow through; W1-3: washes; E1-2: elution fraction; $P_{tot}$: total protein; +Ctl: eGFP:T2A:mCherry clone; F1: total protein extract; F2: thylakoids fragments and mitochondria; F3: plastids; LQ: low quality plastids; MQ: medium quality plastids.

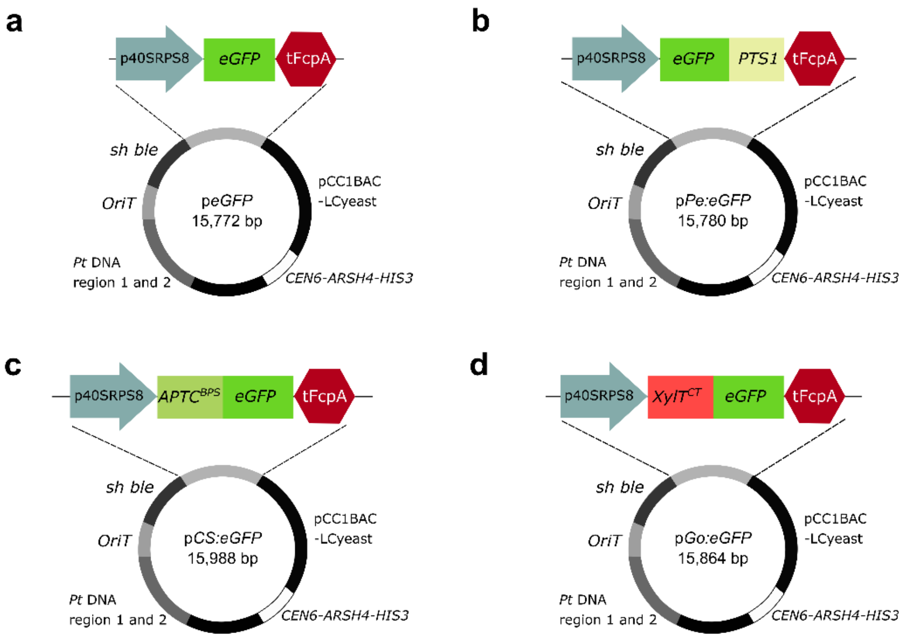


**Figure S3. Constructions of controls for subcellular localization. a**. cytosolic construction *eGFP* without any localization signal; **b**. peroxisome construction harboring the peroxisomal targeting signal 1 (*PST1*); **c**. plastid construction with the bipartite signal of the chloroplast ATPase gamma subunit (*ATPC*$^{BPS}$); **d.** β1,2-xylosyltransferase cytosolic tail and transmembrane domain (CT) for medial Golgi (*XylT*$^{CT}$). *eGFP* expression cassette is antisense from *sh ble*.

**Table S1.** Primer sequences used for Gibson assembly.

| **Name** | **Sequence** |
|---|---|
| FW-BiPCS-Gibson | GCGTTGATCTTGCACCGAAGGAATCAGAGAATAGAATACCA TGAGATCCTTTTGCATCGC |
| RV-BiPCS-Gibson | GTTGCGTGACCGATCGAATGGGGTTATCTTCTTCAAATCCA TCCATGACAATCGTTGCTTTAC |
| FW-SiGol-Gibson | GCGTTGATCTTGCACCGAAGGAATCAGAGAATAGAATACCA TGGCGTTTTTGCCGAATC |
| EV-SiGol-Gibson | GTTGCGTGACCGATCGAATGGGGTTATCTTCTTCAAATCCT GCACTCATCCACATGAGAAA |
| FW-VANCS-Gibson | CTCTGCTCATCGCACGCGTAAAGCAACGATTGTCATGGATG GATTTGAAGAAGATAACCCC |
| FW-VANGol-Gibson | TTTTGCCCTCAATATATGCTTTCTCATGTGGATGAGTGCAGG ATTTGAAGAAGATAACCCC |
| FW-GFPBb-Gibson | TGCCTCGTACCCGATTGTCGCCGTCATTGAAGACGGACGA ATGGTGAGCAAGGGCGAGGAGC |

| | |
|---|---|
| RV-GFPBb-Gibson | CAGCCAAAGTCGAGGTAGTTGTTGCGGTTAGAATTCCTTGTACAGCTCGTCCATGCCGAG |
| RV-GFPPeSi-Gibson | GAAAGTGTCCCAGCCAAAGTCGAGGTAGTTGTTGCGGTTACAACTTCGAGAATTCCTTGTACAGCTCGTCCATGCCGAG |
| FW-BbGFP-Gibson | CTCGGCATGGACGAGCTGTACAAGGAATTCTAACCGCAACAACTACCTC |
| RV-BbGFP-Gibson | GCGTGACCGATCGAATGGGGTTATCTTCTTCAAATCCCATGGTATTCTATTCTCTGATTCCTTCG |
| FW-BbPeSi-Gibson | GACGAGCTGTACAAGGAATTCTCGAAGTTGTAACCGCAACAACTACCTC |
| RV-BbCS-Gibson | ATGCCACAGCCAAAAGGGCTGCGATGCAAAAGGATCTCATGGTATTCTATTCTCTGATTCCTTCG |
| RV-BbER-Gibson | GCAAGGCCAGTGCTGCTACGGCAATTCTCATGAACATCATGGTATTCTATTCTCTGATTCCTTCG |
| RV-BbGol-Gibson | AGGCCACGGTATATCTTCGTCGATTCGGCAAAAACGCCATGGTATTCTATTCTCTGATTCCTTCG |
| 40SRPS8F | GCACCACCCCGGTGAACAGCTCCTCGCCCTTGCTCACCATGGTATTCTATTCTCTGATTCC |
| GFPR | CGTTGATCTTGCACCGAAGGAATCAGAGAATAGAATACCATGGTGAGCAAGGGCGAGG |
| Go40SRPS8F | AGGCCACGGTATATCTTCGTCGATTCGGCAAAAACGCCATGGTATTCTATTCTCTGATTC |
| GoGFPR | GCGTTGATCTTGCACCGAAGGAATCAGAGAATAGAATACCATGGCGTTTTTGCCGAATCG |
| CS40SRPS8F | ATGCCACAGCCAAAAGGGCTGCGATGCAAAAGGATCTCATGGTATTCTATTCTCTGATTC |
| CSGFPR | GCGTTGATCTTGCACCGAAGGAATCAGAGAATAGAATACCATGAGATCCTTTTGCATCGC |
| VANF | GTCGAGGTAGTTGTTGCGGTTAGTGGTGGTGGTGATGGTGTCGTCCGTCTTCAATGACGG |
| PeVANF | GTTGTTGCGGTTACAACTTCGAGTGGTGGTGGTGATGGTGTCGTCCGTCTTCAATGACGG |

**Table S2.** Sequences used in this study.

| Name | Sequence |
|---|---|
| *VpVAN* | GGATTTGAAGAAGATAACCCCATTCGATCGGTCACGCAACGACCCGAT<br>TCCATTGAACCCGCCATTTTGGGCGTTTTGGGTTCGTGCCGCCACGCC<br>TTTCACTTTGCCCGATTTGCCCGCCGATACGGAAAGTCGTACGGATCG<br>GAAGAAGAAATTAAGAAGCGATTTGGAATTTTTGTCGAAAACTTGGCCT<br>TTATTCGATCGACGAACCGTAAGGATTTGTCGTACACGTTGGGAATTAA<br>CCAATTTGCCGATTTGACGTGGGAAGAATTTCGAACGAACCGATTGGG<br>AGCCGCCCAAAACTGTTCCGCCACGGCCCACGGAAACCACCGCTTTG<br>TCGATGGTGTCTTGCCCGTCACGCGTGATTGGCGCGAACAAGGAATTG<br>TATCCCCGGTCAAAGACCAAGGTTCGTGCGGATCGTGCTGGACGTTTT<br>CCACCACGGGAGCCTTGGAAGCCGCCTACACCCAATTGACGGGAAAA<br>TCGACCTCGTTGTCGGAACAACAATTGGTTGACTGCGCTTCGGCCTTT<br>AACAACTTTGGATGTAACGGAGGATTGCCGTCCCAGGCTTTTGAATAC<br>GTCAAATACAACGGTGGAATTGACACCGAACAAACCTACCCCTACCTC<br>GGAGTCAACGGAATTTGTAACTTTAAGCAAGAAAACGTCGGAGTCAAA<br>GTCATTGACTCCATTAACATTACCTTGGGAGCCGAAGATGAATTGAAAC<br>ACGCCGTCGGATTGGTTCGCCCCGTCTCGGTCGCCTTTGAAGTCGTCA<br>AGGGTTTTAACTTGTACAAAAAGGGAGTCTACTCGTCCGACACGTGCG<br>GACGAGATCCAATGGATGTCAACCACGCCGTCTTGGCCGTCGGTTACG<br>GAGTCGAAGACGGTATTCCGTACTGGTTGATCAAAAACTCGTGGGGAA<br>CGAACTGGGGAGATAACGGATACTTTAAGATGGAATTGGGAAAAAACA<br>TGTGCGGAGTCGCCACGTGTGCCTCGTACCCGATTGTCGCCGTCA |
| *eGFP* | ATGGTGAGCAAGGGCGAGGAGCTGTTCACCGGGGTGGTGCCCATCCT<br>GGTCGAGCTGGACGGCGACGTAAACGGCCACAAGTTCAGCGTGTCCG<br>GCGAGGGCGAGGGCGATGCCACCTACGGCAAGCTGACCCTGAAGTTC<br>ATCTGCACCACCGGCAAGCTGCCCGTGCCCTGGCCCACCCTCGTGAC<br>CACCCTGACCTACGGCGTGCAGTGCTTCAGCCGCTACCCCGACCACA<br>TGAAGCAGCACGACTTCTTCAAGTCCGCCATGCCCGAAGGCTACGTCC<br>AGGAGCGCACCATCTTCTTCAAGGACGACGGCAACTACAAGACCCGC<br>GCCGAGGTGAAGTTCGAGGGCGACACCCTGGTGAACCGCATCGAGCT<br>GAAGGGCATCGACTTCAAGGAGGACGGCAACATCCTGGGGCACAAGC<br>TGGAGTACAACTACAACAGCCACAACGTCTATATCATGGCCGACAAGC<br>AGAAGAACGGCATCAAGGTGAACTTCAAGATCCGCCACAACATCGAGG<br>ACGGCAGCGTGCAGCTCGCCGACCACTACCAGCAGAACACCCCCATC<br>GGCGACGGCCCCGTGCTGCTGCCCGACAACCACTACCTGAGCACCCA<br>GTCCGCCCTGAGCAAAGACCCCAACGAGAAGCGCGATCACATGGTCC<br>TGCTGGAGTTCGTGACCGCCGCCGGGATCACTCTCGGCATGGACGAG<br>CTGTACAAGGAATTC |
| *p40SRPS8* | CCCTGCGATAGACCTTTTCCAAACTCACGCAGTCCAAGAAAACAAAGG<br>GGTGAGAAGTATACGCACCTTTCGGTTTCGGCATAATTCTTAAACTCTT<br>GTGGTCACTTTCTTGTGAAGAAGCTAGGGGCACTCGTTTTCCCTCAGA<br>GCCTGCAAACACAAAATTCCTGCAGTCAATTGTCCCAACACTCGGCAA<br>ACCGTATGCGCAAGCAACGATGCGCAGAAGGCCGTGGATGGATGGCG<br>ACTCGCGATATGGCTTCTTGGGTCGCCAGTGTGGTACGTCCGGCGTAT |

| | |
|---|---|
| | GTCAATACGCGAATTCGGACGACTGGCATCTCTAGGAGGAGGATTCCTTCTTTTATGACATGTTTATTTTATATACATTGATGCTTTCCGACAGTCGGAAGTAATAAATGAATTTATTTCAAGACTACCTATACTCCTTTGACTTGTTCGACTAATCTTACCGCTTACTAAAATCTCGAAATCACGCTTGACCTCTCGCACGCAAATTTTTGCTGCTGGACGCTACGCACTCGGCCCAATTCTTCTCGGTCCTCGTCGTCGCAATTGTCGTTGCGTTGATCTTGCACCGAAGGAATCAGAGAATAGAATACC |
| *tFcpA* | CCGCAACAACTACCTCGACTTTGGCTGGGACACTTTCAGTGAGGACAAGAAGCTTCAGAAGCGTGCTATCGAACTCAACCAGGGACGTGCGGCACAAATGGGCATCCTTGCTCTCATGGTGCACGAACAGTTGGGAGTCTCTATCCTTCCTTAAAAATTTAATTTTCATTAGTTGCAGTCACTCCGCTTTGGTTT |
| *ATPC*$^{BPS}$ | ATGAGATCCTTTTGCATCGCAGCCCTTTTGGCTGTGGCATCTGCCTTCACCACACAGCCAACTTCCTTCACTGTGAAGACTGCGAATGTGGGCGAACGGGCGAGTGGGGTTTTCCCTGAGCAGAGCTCTGCTCATCGCACGCGTAAAGCAACGATTGTCATGGAT |
| *PTS1* | TCGAAGTTG |
| *XylT*$^{CT}$ | ATGGCGTTTTTGCCGAATCGACGAAGATATACCGTGGCCTGCTTGTTTATCGGTTTTGCCCTCAATATATGCTTTCTCATGTGGATGAGTGCA |
| *Xa* | TTGAAGACGGACGA |
